# Investigating speech and language impairments in delirium: a preliminary case-control study

**DOI:** 10.1101/351007

**Authors:** Samantha Green, Satu Reivonen, Lisa-Marie Rutter, Eva Nouzova, Nikki Duncan, Caoimhe Clarke, Alasdair M. J. MacLullich, Zoë Tieges

## Abstract

**Introduction**

Language impairment is recognized as a diagnostic feature of delirium, yet there is little neuropsychological research on the nature of this dysfunction. Here we hypothesized that patients with delirium show impairments in language formation, coherence and comprehension.

**Methods**

This was a case-control study in 45 hospitalized patients (aged 65-97 years) with delirium, dementia without delirium, or no cognitive impairment (N=15 per group). DSM-5 criteria were used for delirium. Speech was elicited during (1) structured conversational questioning, and (2) the “Cookie Theft” picture description task. Language comprehension was assessed through standardized verbal and written commands. Interviews were audio-recorded and transcribed.

**Results**

Delirium and dementia groups scored lower on the conversational assessment than the control group (p<0.01, moderate effect sizes (r) of 0.48 and 0.51, resp.). In the Cookie Theft task, the average length of utterances (i.e. unit of speech), indicating language productivity and fluency, distinguished patients with delirium from those with dementia (p<0.01, r=0.50) and no cognitive impairment (p<0.01, r=0.55). Patients with delirium performed worse on written comprehension tests compared to cognitively unimpaired patients (p<0.01, r=0.63), but not compared to the dementia group.

**Conclusions**

Production of spontaneous speech, word quantity, speech content and verbal and written language comprehension are impaired in delirious patients compared to cognitively unimpaired patients. Additionally, patients with delirium produced significantly less fluent speech than those with dementia. These findings have implications for how speech and language are evaluated in delirium assessments, and also for communication with patients with delirium.

## Introduction

Delirium is a severe neuropsychiatric syndrome characterized by acute disturbances in attentional functioning and a range of other cognitive deficits and neuropsychiatric symptoms [1-3]. Language dysfunction is explicitly included within the DSM-5 criteria for delirium (under Criterion C: other cognitive disturbance [4]). However, the DSM guidance notes do not specify which domains of language are affected and how these language deficits should be measured. Language dysfunction is listed in several delirium rating scales (Table 1) but most do not include explicit evaluation of language, with the exception of the Delirium Rating Scale Revised-98 (DRS-R98 [5]).

**Table 1.** Language disturbances in delirium diagnostic tools.

| Assessment tool | Cognitive feature | Description/Test of language impairment |
| --- | --- | --- |
| Delirium Rating Scale-Revised-98 [5] | Language disturbance | 0=Normal language<br>1=Mild impairment including word-finding difficulty or problems with naming or fluency<br>2=Moderate impairment including comprehension difficulties or deficits in meaningful communication (semantic content)<br>3=Severe impairment including nonsensical semantic content, word salad, muteness, or severely reduced comprehension |
| 3D-Confusion Assessment Method [13] | Disorganized thinking | Observational measure: Conversation rambling, off target, or abnormally sparse |
| Confusion Assessment Method [14] | Disorganized thinking | Rambling or irrelevant conversation, unclear or illogical flow of ideas, or unpredictable switching from subject to subject |
| Cognitive Test for Delirium [15] | Comprehension | Will a stone float on water? / Will a leaf float on water?<br>Can you use a hammer to pound nails? / Is a hammer good for cutting wood?<br>Do two pounds of flour weight more than one? / Is one pound heavier than two? |
|  |  | Will water go through a good pair of rubber boots? / Will a good pair of rubber boots keep water out? |
| Delirium Observation<br>Screening Scale [16] | Thinking | Gives answers that do not fit the question<br>Talks slowly or answers slowly<br>Reacts slowly to instructions<br>Speaks incoherently |
| Delirium Symptom<br>Interview [17] | Incoherent speech | Was the patient's speech:<br>a) Unusually limited or sparse (e.g. yes/no answers)<br>b) Unusually slow or halting<br>c) Unusually slurred<br>d) Unusually fast or pressured<br>e) Unusually loud<br>f) Unusually repetitive (e.g. repeats phrase over and over<br>g) Have speech sounds in the wrong place<br>h) Have words or phrases that were disjointed or inappropriate<br>(Questions a-h scored on a scale of 1[no]-4 [severe]) |
|  |  | i) If present did the patient's speech fluctuate during the interview, for example, patient spoke normally for a while, then sped up (yes / no) |
| Delirium-O-Meter [18] | Incoherence | 0= What the patient says is easy to understand even for someone who does not know him very well<br>1=What the patient says is not always easy to understand, sometimes jumps from one topic to another<br>2=Clearly hard to follow, associative, sentences appear unrelated, sometimes stops in the middle of a sentence<br>3=Not able to express a coherent thought, unfinished sentences, loose words, yells, moaning |
| Delirium Motor Subtyping Scale [19] | Indicator of hypoactive delirium | Decreased amount of speech evidenced by a positive response to either:<br>Does (s)he speak less than before?<br>Is (s)he lacking in spontaneous speech<br>? E.g. only speaks when spoken to. |

Studies on communication difficulties in delirious patients have reported significant issues when conveying information, as reported by patients, their relatives and nursing staff [6-8]. Yet very few studies have investigated the nature of language abnormalities in delirium. Fundamental domains of language such as speech production in delirium have remained largely unexplored despite one study reporting language impairments in over half of patients with delirium [1].

Wallesch and Hundsalz [9] compared the performance on single word naming and comprehension tasks between patients with delirium and those with Alzheimer’s dementia. Delirious patients produced more perseverations and semantically unrelated misnamings compared to the dementia group. Another study analyzed writing disturbance (dysgraphia), reporting that signature writing was impaired in delirium [10]. Additionally, the production of jagged and angular segments of letters was found to be specifically impaired in delirium compared with non-delirious psychiatric inpatients [11]. Chedru and Geschwind [12] compared writing abilities of hospitalized patients with and without delirium. Dysgraphia was almost always present in delirium, typically involving motor and spatial aspects of writing,and spelling and punctuation errors [12]. Tate and colleagues [20] explored the communication of symptoms between critically ill patients and nurses, finding that patients with delirium were less likely to initiate symptom communication compared with non-delirious patients.

The present study aimed to investigate language production and comprehension in delirium. We hypothesized that patients with delirium would produce fewer words and more errors in language production. Specifically, we predicted that patients with delirium would produce more irrelevant speech content and more unrelated or inappropriate content elements on language production tasks than patients with dementia (without delirium) or no known cognitive impairment. We also hypothesized that language comprehension would be poorer in groups with delirium compared to dementia or no cognitive impairment, with more comprehension-related errors in delirium (as reported by [9]).

## Methods

### Design

This was a case-control study including three groups of hospitalized patients: (1) patients with delirium (with or without dementia), (2) patients with a formal diagnosis of dementia but not current delirium, and (3) patients without cognitive impairment. The study obtained a favorable opinion from the Scotland A Research Ethics Committee.

### Participants

Patients were recruited from the orthopedics and Medicine of the Elderly wards at the Royal Infirmary of Edinburgh, Scotland. Patients aged 65 years and older, fluent in English and able to provide written informed consent, or those with a suitable proxy, were eligible to participate. Exclusion criteria were: sensory impairment severe enough to hinder cognitive testing, severe illness where clinical staff considered study participation to pose a risk to patient care, or a history of dysarthria, aphasia, and traumatic brain injury.

Two researchers (SG and SR) approached a total of 100 individuals each identified by the clinical team as potentially eligible for the study. Of these, 25 patients declined to participate, 13 were excluded because either the proxy declined or could not be contacted, seven were ineligible and grouping for the remaining 10 patients was undetermined (i.e. they did not fit any of the pre-specified clinical groups). The final study sample contained 45 participants (N=15 per group; 26 females).

## Measures and Procedures

### Reference standard assessment

Cognitive and delirium assessments were conducted independently by two psychology graduates (SG and SR). The researchers had been trained by a senior geriatrician (AM), a psychology research fellow (ZT) and two researchers (EN and LMR), one of them (LMR) a registered speech and language therapist. The researcher who identified patients and obtained consent completed the reference standard assessment which tested different domains of cognition including memory, attention and orientation. This assessment lasted approximately 20 minutes and results were used alongside observational and medical information to determine group allocation. The following assessment tools were used:

1. A brief memory test in which patients were shown drawings of a lemon, a key and a ball and asked to repeat all three items immediately (short-term recall) and at the end of the assessment (5-10 min delay; long-term recall) [21].
2. The Short Orientation-Memory-Concentration test (OMCT), a six-item cognitive test predominantly focused on measuring orientation and working memory [22]. The maximum possible OMCT score was 28, with scores of 20 or below indicating cognitive impairment.
3. A brief attentional test battery comprising digit span forwards and backwards, and days of the week and months of the year backwards [23]. The maximum possible score was 7, with a score of 5 or below suggesting attentional impairment. The Vigilance A task from the Montreal Cognitive Assessment [24] was administered as an additional measure of sustained and focused attention, whereby one error was permitted.
4. The UVA Pain Rating Scale (online available at https://uvahealth.com/patients-visitors/images/documents/UVAPainRatingScale.pdf (Accessed on 18 January 2016)), a numerical pain rating scale supplemented with a faces pain thermometer (adapted from [25]) was used to assess patients’ pain intensity. Scores ranged from 0 (no pain) to 10 (worst pain imaginable).
5. The Delirium Rating Scale-Revised-98 (DRS-R98 [5]) was used to aid delirium assessment, supplemented by the Observational Scale of Level of Arousal (OSLA [26] and the Richmond Agitation-Sedation Scale (RASS; [27, 28]) for measuring level of arousal.

### Delirium diagnosis according to DSM-5

Delirium was ascertained using DSM-5 diagnostic criteria. Specifically, cognitive scores obtained during the reference standard assessment, observations of the patient’s behavior (captured in OSLA, RASS and DRS-R98 items) and information obtained from informants, records on pre-admission functional status and case notes were all considered for the delirium diagnosis. Where group assignment was uncertain, cases were discussed amongst the study team and consensus reached by AM and ZT, blind to the results of the speech and language assessment. Additionally, known dementia diagnosis in the delirium group was ascertained by case notes and/or discussions with the medical team.

### Linguistic task battery

A short battery of language tasks was completed within one hour of completing the reference standard assessment to minimize the impact of delirium’s fluctuating nature. The researcher administering the language assessment was blinded to the participant’s group allocation and any other information obtained during the reference standard assessment.

The following domains of language were assessed:

#### 1. Language production

##### 1a. Conversational speech

Firstly, a conversationally based test of spontaneous speech was administered by adapting three questions from the Western Aphasia Battery-Revised [29]. The researcher transcribed patient responses and separated speech into utterances (units of speech, as previously defined by [30]). Utterances could be fragmented or complete sentences, separated by a significant pause. The total number of utterances, and the number of relevant and irrelevant utterances was determined and the percentage of relevant utterances was calculated (see Table 2 for details). The resulting conversational speech score ranged from 0-9 with higher scores indicating more relevant speech content.

**Table 2.**
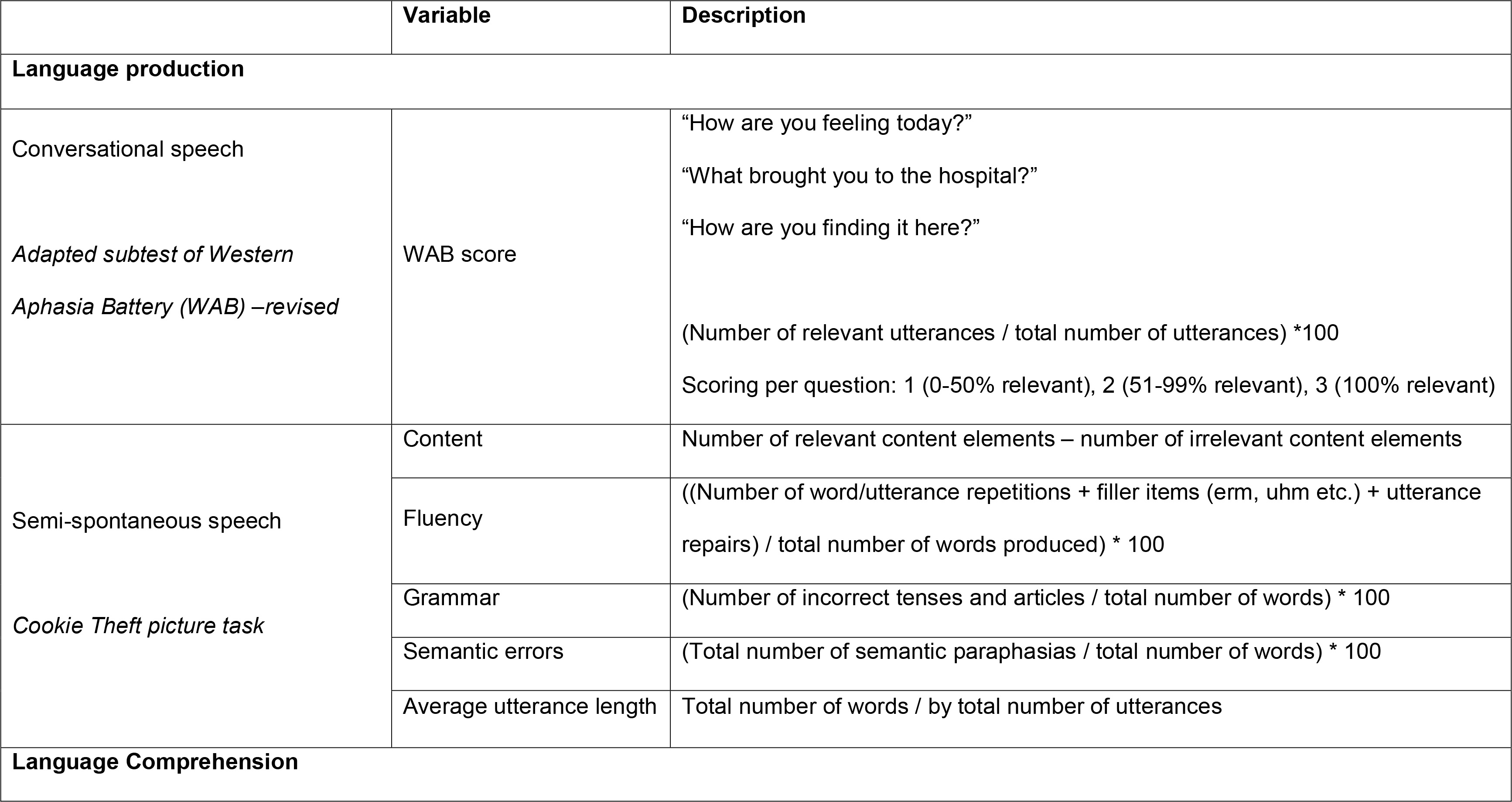
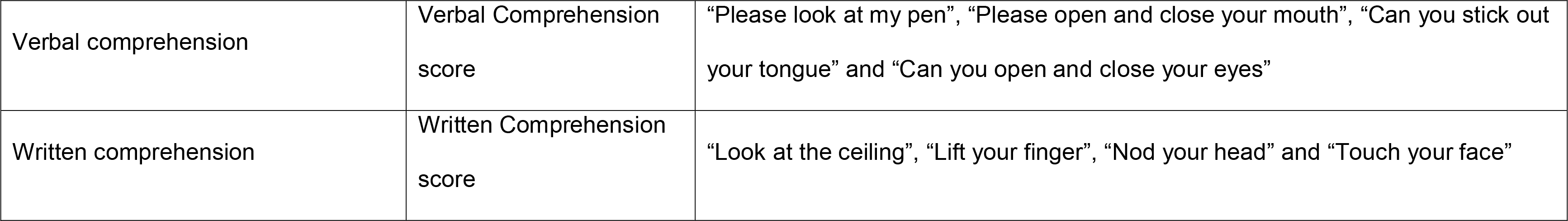
Language assessment: task variables derived from the conversational speech assessment and Cookie Theft picture task.

##### 1b. Semi-spontaneous speech

Semi-spontaneous speech was elicited using the Cookie-Theft Picture Description Task from the Boston Diagnostic Aphasia Examination ([31], a widely used test of language production which has been used in healthy adults ([32] and patients including those with Alzheimer’s dementia [33]. Participants were asked to describe what they saw in the picture and prompted once with ‘do you see anything else?’. The researchers transcribed descriptions of the image and determined (i) the total number of words and utterances and (ii) the average utterance length to provide an indicator of overall language production and fluency (Table 2). Speech samples were then examined for four elements: speech content, speech fluency, grammar and semantic paraphasias [34].

###### Speech content

Speech content was divided into two categories, ‘relevant content’ or ‘irrelevant content’. An overall ‘content elements’ score was calculated, reflecting the number of relevant concent elements minus irelevant items (Table 2).

###### Speech fluency

Issues with fluency were identified within transcripts if they fell into one of the following categories: word or utterance repetition, filler items (in accordance with [35] e.g. ‘erm’), and utterance repairs where the speaker goes back and changes something he or she just said. An overall fluency score was calculated, reflecting the number of errors in relation to the total amount of speech produced (Table 2).

###### Grammar

Grammar errors were accounted for by summing any incorrect tenses and/or the use of ambiguous articles and captured in an oveall grammar score (Table 2). Ambiguous articles refer to the use of a word such as ‘that, this, it,’ etc. in which the patient does not name an object or person.

###### Semantic paraphasias

Semantic paraphasias, referring to words of a similar or unrelated category which are used to describe an object (e.g. the word chair instead of stool [36]) were captured in an overall semantic error score (Table 2).

Once scoring was completed, a subset of 15 (5 from each clinical group) transcripts of the Cookie Theft Picture Description Task was sent to two trained researchers (ND and CC) who were blinded to participant grouping to provide inter-rater scores.

##### 2. Language comprehension

Participants were presented with four verbal and four written commands, printed on A4 paper printed in 130 point Calibri upper case font (Table 2). One point was given for each action performed correctly. Verbal and written comprehension scores were recorded separately.

### Transcription and statistical analysis

Participants’ responses to the speech tasks were recorded with an Olympus VN-5500PC audio recorder and transcribed verbatim using conventions (adapted from [37]).

Non-parametric tests were used due to the high prevalence of non-normally distributed data according to the Shapiro-Wilk test of normality. Kruskall-Wallis tests were used to explore differences between groups (delirium, dementia, no cognitive impairment). Mann-Whitney U tests were used for the pairwise comparisons. Cohen’s effect sizes (r) were reported for significant effects on the language variables. Associations between level of arousal and attention performance on language ability was assessed using Kendall’s tau-b rank correlation coefficients as there were ties in the data. Cohen’s effect sizes were used when interpreting correlation coefficients and strength, thus correlations >0.5 were classified as ‘large’ (Cohen, Cohen, West & Aiken, 2003). Inter-rater reliability was evaluated using intra-class correlations. Statistical analyses were completed in IBM SPSS Statistics 22.0.0.1. Statistical significance was considered as a two-sided p value <0.01.

## Results

### Inter-rater agreement

Inter-rater agreements for the Cookie Theft picture description task scoring were satisfactory, with intra-class correlation coefficients ranging from 0.63 to 0.99 [38].

### Participants

Patients with delirium and dementia were overall older than controls (delirium vs. control: U= 23.00, p<0.001; dementia vs. control: U=49.5, p<0.01). Age did not differ between dementia and delirium groups. There was no difference in sex between groups. The cognitive tests used in the group assessments showed differences commensurate with group allocations. OMCT scores were higher in cognitively unimpaired patients indicating better overall cognition (median=26, Inter-Quartile Range (IQR)=26-28) compared to patients with delirium (median=0, IQR=0-6; U=0.00, p<0.001) and dementia (median=4, IQR=3-6; U=2.00, p<0.001; Table 3). Delirium and dementia groups did not differ on performance on the cognitive tests. Nine patients with delirium had a diagnosis of dementia (3 Alzheimer’s dementia, 1 mixed dementia, 5 unspecified).

**Table 3.**
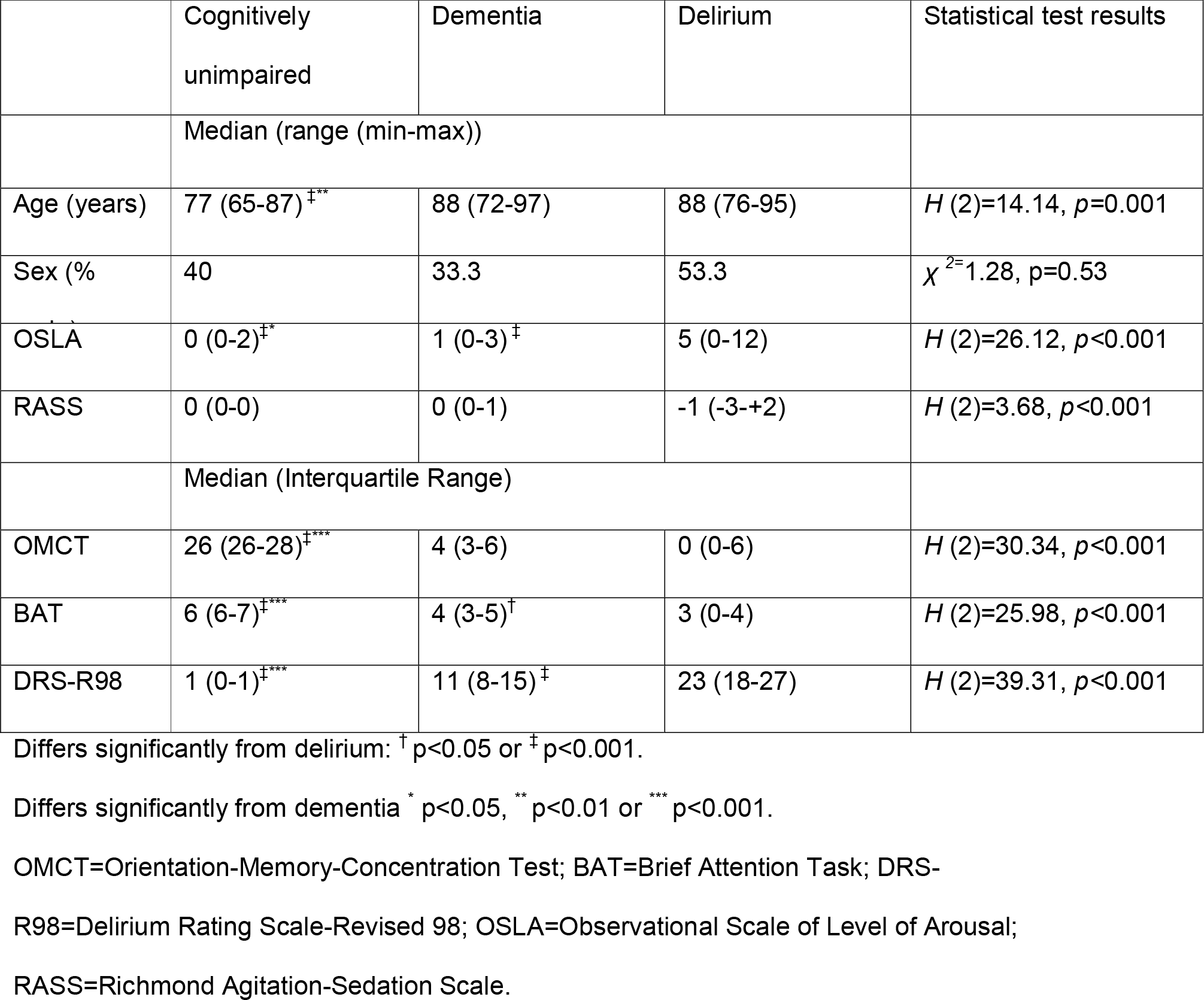
Descriptive statistics for cognitive tests and behavioral scales.

## Group differences in language production and comprehension

### Language production

Performance on the conversational speech assessment differed among groups (H=9.57, p <0.01). Delirium and dementia groups produced speech that was less relevant to the questions (delirium: median=7, IQR=5-8; dementia: median=7, IQR=5-8) compared to the cognitively unimpaired group (median=9, IQR=8-9; delirium vs. control: U=52.50, p <0.01, r=0.48; dementia vs. control: U=48.00, p<0.01, r=0.51). There were no differences between delirium and dementia groups (U=107.00, p=0.82; Fig 1 and S1 Table).

**Fig. 1.**
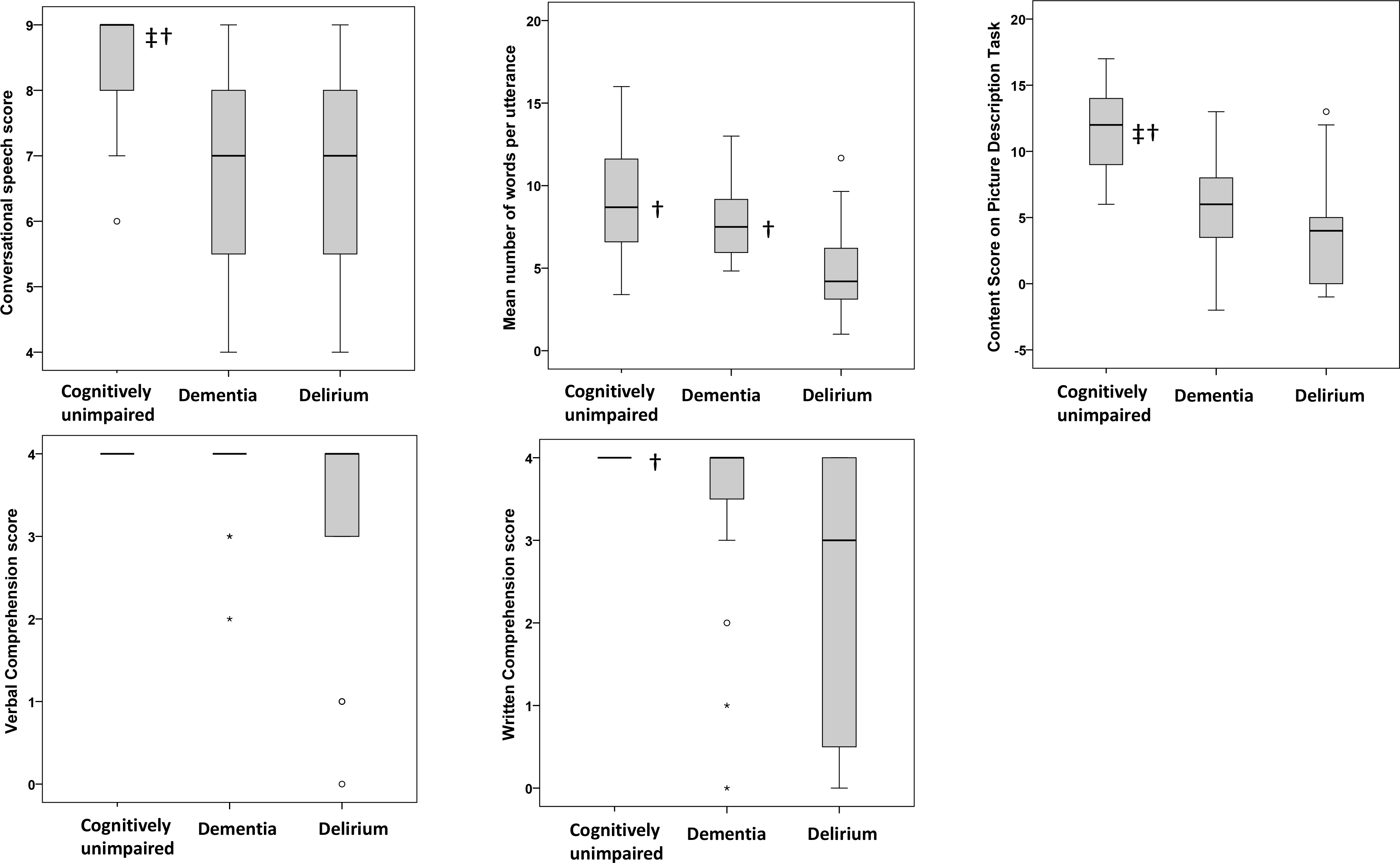
Group results for the conversational speech assessment (A), the average number of words per utterance (Cookie Theft picture task) (B), the content score (Cookie theft) (C), verbal comprehension score (D) and written comprehension score (E). The interquartile range and median value of each dataset are represented by the height of the inner box and the position of the central horizontal line, respectively. The positions of the upper and lower bars of each plot indicate the maximum and minimum non-outlier values of each dataset. Any outliers are represented by open circles on the plot. Symbols to the right of the median lines indicate group scores that differ significantly from delirium (†) or dementia (‡).

Furthermore, groups differed in performance on the Cookie Theft Picture Description Task. Firstly, the average number of words per utterance varied between groups (H=11.62, p<0.01). Specifically, patients with delirium produced fewer words (median=4.2, IQR=3-7) than those with dementia (median=7.5, IQR=5.43-9.33; U= 47.00, p <0.01, r=0.50) and cognitively unimpaired patients (median=8.69 words, IQR=6.33-11.82; U=40.50, p<0.01, r=0.61).

Regarding the content, fluency, grammar and semantics measures, only the content score differed between groups (H=17.74, p<0.001). Cognitively unimpaired patients produced more relevant content (median=12, IQR=9-14) than those with delirium (median=4, IQR=0-5; U=23.50, p<0.001, r=0.68) and dementia (median=6, IQR=3-7, U=33.00, p<0.01, r=0.61). There were no differences between dementia and delirium groups (U=78.00, p=0.15).

### Language comprehension

There were no differences between groups in verbal comprehension (H=7.57, p=0.02), but groups differed in written comprehension (H=12.54, p<0.01). This was poorer in delirium (median=3, IQR=0-4) compared to cognitively unimpaired patients (median=4, IQR=4-4; U=45.00, p<0.01, r=0.63). Delirium and dementia patients did not differ (dementia: median=4, IQR=3-4; U=74.5, p=0.08). There was no difference between dementia and cognitively unimpaired groups (U=82.50, p=0.04).

### Associations between measures of attention, arousal and language

Moderate-to-strong associations were found between scores on the brief attentional test scores and the total number of words per utterance in the Cookie Theft task (тb=0.43, p<0.01), speech content (тb=0.60, p<0.001), verbal comprehension (тb=0.59, p<0.001) and written comprehension (тb=0.62, p<0.001) in the sample as a whole, reflecting more effective language production and comprehension with better attentional function.

Higher OSLA arousal scores (indicating greater abnormality) were moderately-well correlated with less conversational speech (тb=-0.31, p<0.01), fewer words per utterance (тb=-0.34, p<0.01), reduced semantic content (тb=-0.44, p<0.01) and more fluent speech content (тb=0.30, p<0.01). Higher absolute RASS scores (indicating greater abnormality) were associated with fewer words per utterance (тb=-0.44, p<0.01), reduced semantic content (тb=-0.52, p<0.01), and verbal (тb=-0.51, p<0.01) and written (тb=-0.51, p<0.01) comprehension.

## Discussion

The novel findings in this study are that patients with delirium showed multiple abnormalities in speech and language production and comprehension compared with cognitively unimpaired patients. Hospitalized patients with dementia also displayed multiple abnormalities, specifically in language production (utterance length and speech content) but not comprehension. With respect to language production, individuals with delirium produced more irrelevant speech than cognitively unimpaired participants. Additionally, patients with delirium produced significantly shorter utterances in the Cookie Theft task than patients with dementia or those without cognitive impairment, and identified less relevant and more irrelevant content items than both comparison groups. These findings support our hypotheses and suggest that the delirium group had difficulties producing relevant speech content, even when prompted by a visual stimulus. There were no differences in overall fluency, grammar and semantic scores derived from the Cookie Theft task (though the finding of shorter utterances in delirious patients does suggest impaired fluency in this group). Verbal language comprehension did not differ between groups, but individuals with delirium or dementia had worse *written* comprehension compared to controls.

The moderate-to-strong associations between arousal and attention with language performance support the notion that language dysfunction in delirium and dementia may in part be secondary to more fundamental disturbance in arousal and cognition. This possibility has been suggested in relation to another common feature of delirium, disorganised thinking [2]. However the variation in the presence and severity of the language abnormalities in the delirium group in the present study, and the variable relationship of language abnormalities with arousal and cognition within this group suggest that they may also be a distinct neuropsychological disruption in delirium.

Language and communication difficulties are a recognized part of the delirium syndrome [1, 6-8] but have been subject to very little systematic investigation. Previous studies have reported impairments in word naming, comprehension and handwriting disturbance in delirium [9, 10, 12]. The current paper expands upon this work through systematic investigation of verbal language production via conversational speech and the Cookie Theft picture description task as well as basic language comprehension in delirium and dementia. To our knowledge this is the first study to demonstrate feasibility of using a well-validated picture description paradigm to prompt speech in patients with delirium. Adopting such systematic approaches to language assessment are needed to advance our understanding of language dysfunction as part of the neuropsychological profile of delirium. Importantly, we used a multidimensional linguistic approach in the assessment by capturing multiple aspects of language including comprehension, grammar, speech content and speech fluency. This holistic approach generates patholinguistic profiles which are clinically more relevant than studying one single domain or aspect of language, reflecting “real life” communication or language discourse.

The impairments in language production and comprehension identified in this study have practical and theoretical implications. Firstly, with regard to communicating with delirious patients in clinical settings, the range of impairments seen in delirium highlight the need for communication strategies adapted to the respective needs of patients and delirium-focused communication guidelines. It is prudent that clinical staff briefly assess a patient’s ability to produce and comprehend language prior to engaging in conversations about treatment options, informed consent, and so on. Efficient communication between clinician and patient is also essential for effective management of pain and dehydration, which are a common concern in people with delirium. Taken together, our study findings suggest that clinical staff should be sensitized for communication disturbances in delirious patients, similar to how this is implemented for patients with dementia or primary aphasic disorders. Given the substantial difficulties in written comprehension observed in patients with delirium as well as dementia in study, particular attention should be paid to determining if such patients have understood written materials.

A second implication relates to the utility of language assessment in diagnosing delirium. The present findings suggest that language observations during routine interview and cognitive testing may contribute usefully to the diagnostic process. However, although language impairments are clearly present and prominent in delirium, our study does not suggest that delirium can be distinguished from dementia based on language impairments alone. Specifically, findings from the conversational task suggest that content of speech, when asked how the patient was feeling and their reason for hospitalization, could not discriminate delirium from dementia. Thus, clinicians may not be able to identify delirium in older people merely through informal conversations with them. Rather, asking questions which require more complex language abilities, aided by visual prompts to help elicit speech, may be a more effective means of testing for delirium than simply conversing with the patient.

Nonetheless, because not all delirium occurs in patients with dementia, the value of speech and language testing as part of delirium diagnosis, for gauging severity of delirium and for monitoring recovery of delirium, is of interest and requires further investigation. Brief language production and comprehension tasks have the potential to add usefully to the delirium screening or diagnostic process.

The limitations of this study include the case-control design (selective study sample) and relatively small patient numbers. The language battery was adapted solely for this study and has not been validated previously in delirium. Some patients may have had undiagnosed dementia or cognitive impairment. In future studies with larger samples, it would be desirable to examine deficits in patients with known dementia status and also in patients with delirium but without pre-existing cognitive impairment. Finally, the small sample size did not permit consideration of specific language profiles associated with hypo-and hyperactive and mixed delirium subtypes [19].

In spite of the limitations, given the finding that reduced utterance length was characteristic of delirium, it would be useful to further operationalize this variable. Future work could examine quantifiers of language as indicators of delirium, with the potential to assist in the diagnosis of sub-syndromal forms often missed, and also potentially the monitoring of delirium severity over time. Further, other aspects of language such as word finding and pronunciation could be explored. In larger samples, variables such as age, socioeconomic status and geographical location would need to be accounted for as these are known to influence language performance (as discussed in [39]). It would also be useful to explore language abilities in patients with delirium but without dementia (for example in a critical care population) and to identify whether language disturbances could be used as a measure of delirium severity and recovery from delirium.

The present study supports and expands on previous findings of language impairment in delirium ([10-12]. The findings provide additional characterization of language disturbances in delirium, which though present in in DSM-5 are not specified in detail. Further, these results provide a basis for future research on language abnormalities in delirium which could lead to improve the identification, assessment and monitoring of this serious disorder. Clinically, the present study suggests the need for increased awareness of language production and comprehension difficulties in delirium and delirium-specific guidance, since effective patient-clinician communication is central to good clinical care.

## Acknowledgments

The authors thank patients and staff from the Medicine of the Elderly and acute orthopedic wards of the Royal Infirmary of Edinburgh.

## Supporting information

**S1 Table. Descriptive statistics and statistical group comparisons for scores on the language assessment**. IQR = Inter-Quartile Range. Variables derived from the Cookie theft picture description task are denoted with a *. Kruskal-Wallis and Mann-Whitney U were used.

